# Structural Basis for Selective Proteolysis of ADAM10 Substrates at Membrane-Proximal Sites

**DOI:** 10.1101/2022.10.22.513345

**Authors:** Colin H. Lipper, Emily D. Egan, Khal-Hentz Gabriel, Stephen C. Blacklow

**Affiliations:** Department of Biological Chemistry and Molecular Pharmacology, Blavatnik Institute, Harvard Medical School, Boston, MA 02115, USA; Selecta Biosciences, Watertown, MA 02472 USA; Department of Cancer Biology, Dana Farber Cancer Institute, Boston, MA 02215, USA

## Abstract

The endopeptidase ADAM10 is a critical catalyst for regulated proteolysis of key drivers of mammalian development and physiology, and for non-amyloidogenic cleavage of the Alzheimer’s precursor protein as the primary α-secretase. ADAM10 function *in vivo* requires formation of a complex with a C8-tetraspanin protein, with different ADAM10-C8-tetraspanin complexes having distinct substrate selectivity, yet the basis for such selectivity remains elusive. We present here a cryo-EM structure of a vFab-ADAM10-Tspan15 complex, which shows that Tspan15 binding relieves ADAM10 autoinhibition and positions the enzyme active site about 20 Å from the plasma membrane for membrane-proximal substrate cleavage. Cell-based assays of N-cadherin shedding establish that the positioning of the active site by the interface between the ADAM10 catalytic domain and the bound tetraspanin influences selection of the preferred cleavage site. Together, these studies reveal the molecular mechanism underlying selective ADAM10 proteolysis at membrane-proximal sites and offer a roadmap for its modulation in disease.

## Introduction

Ectodomain shedding of transmembrane proteins plays a central role in a wide range of normal and pathophysiologic processes (Lichtenthaler et al., 2018; Peschon et al., 1998). A Disintegrin and Metalloproteinase 10 (ADAM10) is a single-pass transmembrane protease that is essential in mammals (Hartmann et al., 2002), cleaving its substrates at membrane-proximal extracellular sites in a process called ectodomain shedding (Kuhn et al., 2016; Lambrecht et al., 2018; Pruessmeyer and Ludwig, 2009; Weskamp et al., 2006). ADAM10 functions in physiological Notch signaling by catalyzing ligand-dependent Notch activation during development (Mumm et al., 2000; Sprinzak and Blacklow, 2021), and in pathologic Notch signaling in T-cell acute lymphoblastic leukemia (Weng et al., 2004), where oncogenic mutations result in dysregulated cleavage (Sulis et al., 2011). Additionally, ADAM10 processes a number of adhesion proteins including various cadherins, where cleavage is associated with both physiological and tumor cell migration (Kohutek et al., 2009; Maretzky et al., 2005; Solanas et al., 2011), and Neuroligin-3, the shedding of which is indispensable for growth of high-grade gliomas (Venkatesh et al., 2017). ADAM10 also acts as an alpha-secretase in cleaving the amyloid precursor protein (APP) in a non-amyloidogenic manner, preventing toxic amyloid-beta generation by beta- and gamma-secretase cleavage (Esler and Wolfe, 2001; Kuhn et al., 2010). Moreover, loss-of-function mutations of ADAM10 are associated with an increased risk of developing Alzheimer’s disease (Suh et al., 2013).

ADAM10 is produced as an inactive zymogen, which matures into the active enzyme upon proprotein convertase-catalyzed release of its prodomain in the trans-Golgi network (Anders et al., 2001). The structure of the mature ectodomain, which includes metalloproteinase, disintegrin, and cysteine-rich domains, revealed an autoinhibited conformation in which the cysteine-rich domain contacts the metalloproteinase domain, partially occluding access to the active site (Seegar et al., 2017).

The maturation and function of ADAM10 are modulated by a class of proteins called tetraspanins, four-pass integral membrane proteins that have a vital role in organizing protein complexes in the membrane (Hemler, 2005). The six members of the TspanC8 subfamily (Tspan5, Tspan10, Tspan14, Tspan15, Tspan17, Tspan33; so named for their eight extracellular cysteines) are essential regulators of ADAM10 (Harrison et al., 2021), required for its maturation, trafficking, and subcellular localization (Dornier et al., 2012; Haining et al., 2012; Harrison et al., 2021; Shah et al., 2018). Several studies also indicate that TspanC8 proteins can dictate ADAM10 substrate specificity (Dornier et al., 2012; Eschenbrenner et al., 2020; Jouannet et al., 2016; Lipper et al., 2022; Noy et al., 2016a; Prox et al., 2012; Seipold et al., 2018). For example, Tspan15 is the only TspanC8 protein that promotes cleavage of N-cadherin (Ncad) (Noy et al., 2016b), whereas Tspan5 and Tspan14 both favor cleavage of Notch receptors (Dornier et al., 2012). Tspan15-dependent N-cadherin cleavage is especially relevant to cancer progression and is associated with tumor cell invasion and metastasis (Hiroshima et al., 2019; Kohutek et al., 2009; Lipper et al., 2022; Zhang et al., 2018), highlighting the role of different Tspans as extrinsic factors in regulating ADAM10 cleavage selectivity, distinct from the intrinsic regulation imposed by selective binding pockets on the protease itself.

To elucidate how TspanC8 proteins recognize ADAM10 and understand how complex formation positions the ADAM10 active site for substrate cleavage, we determined the structure of a complex between Tspan15 and ADAM10 by cryo-EM, revealing an open conformation of ADAM10 positioned by the bound Tspan15 for efficient cleavage of transmembrane substrates at membrane-proximal positions. Our structural analysis and cell-based assays reveal both intrinsic and extrinsic regulatory mechanisms for ADAM10 catalyzed proteolysis of native substrates, and show that the identity of the bound TspanC8 partner protein can tune substrate specificity by altering the position of the ADAM10 active site relative to the plasma membrane.

## Results

### Structure of the ADAM10-Tspan15 complex

To facilitate structure determination, we first co-expressed ADAM10 and Tspan15 in HEK293F cells and extracted the ADAM10-Tspan15 complex into n-Dodecyl-β-D-Maltoside (DDM) – cholesterol hemisuccinate (CHS) detergent micelles. After affinity purification and exchange into glyco-diosgenin (GDN) – CHS, we then added an anti-ADAM10 11G2 Fab fragment, which binds to the disintegrin domain of ADAM10 without affecting its conformation (Seegar et al., 2017), as a fiducial marker for cryo-EM analysis and to increase complex mass, and included the metalloproteinase inhibitor BB94 during purification to prevent sample degradation from ADAM10 proteolytic activity. After isolating the full 11G2 Fab-ADAM10-Tspan15 complex by size-exclusion chromatography (Figure S1A-C, related to Figure 1) and confirming that the ADAM10-Tspan15 complex was catalytically active (Figure S1D, related to Figure 1), we determined the structure of the vFab-ADAM10-Tspan15 complex to 3.3 Å resolution using cryo-EM (Figure S2, related to Figure 1).

**Figure 1.**
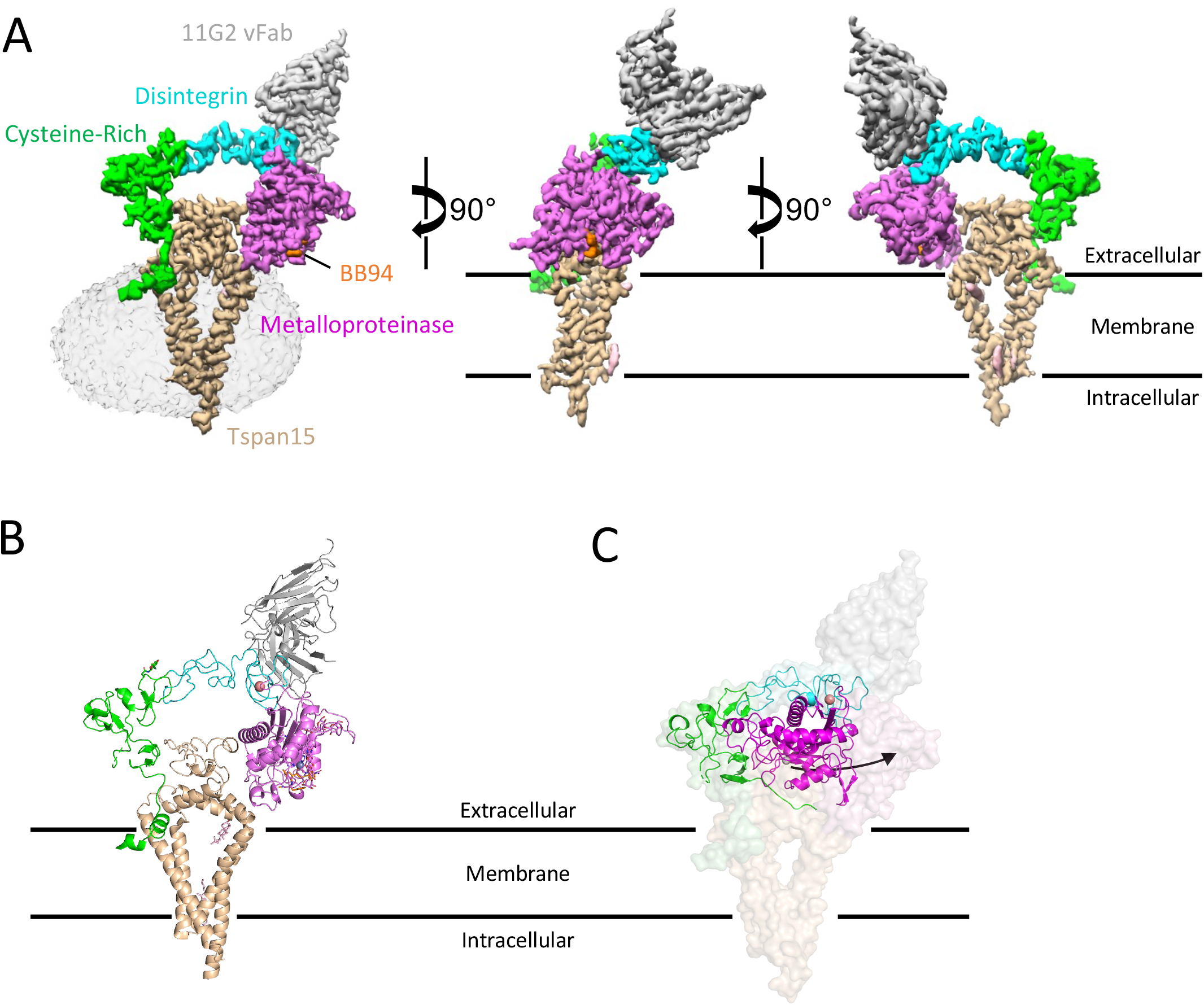
Structure of the ADAM10-Tspan15 complex. **A.** Different views of the cryo-EM electron density map of the vFab-ADAM10-Tspan15 complex. The density associated with the detergent micelle is rendered transparently in the left panel. Tspan15 is beige, the 11G2 vFab is gray, the BB94 inhibitor is orange, and the catalytic, disintegrin, and cysteine-rich domains of ADAM10 are magenta, cyan and green, respectively. **B.** Cartoon representation of the complex, using the same color scheme. The zinc ion at the active site is represented as a gray sphere, and the bound calcium ion in the disintegrin domain is orange. **C.** Comparison of the free ADAM10 ectodomain (ribbon) with ADAM10 in the vFab-ADAM10-Tspan15 complex (surface). See also Figures S1–S3, Table S1, and Supplementary Video S1.

To build the initial model, the coordinates of the crystal structure of the ADAM10 ectodomain bound to the 11G2 Fab (Seegar et al., 2017) and the alphafold2 model for Tspan15 (Jumper et al., 2021) were docked into the cryo-EM electron density map. Whereas the Tspan15 model and the vFab portion of 11G2 fit readily into the density map, the ectodomain of ADAM10 did not. Instead, the catalytic domain and the disintegrin + cysteine-rich domains of ADAM10 were docked separately into the electron density before further refinement to produce a final model.

In the complex, the ADAM10 ectodomain adopts a C-shaped conformation encircling the Tspan15 ectodomain, and the two ectodomains are nestled together atop the four Tspan15 intramembrane helices which adopt a conical arrangement (Figure 1A,B). C-terminal to I670 of ADAM10, the transmembrane and cytoplasmic regions are not visible even when bound, presumably because they remain disordered in the complex. The variable domain of the 11G2 Fab (vFab) caps the complex at the N-terminal end of the ADAM10 distintegrin domain, distant from sites of ADAM10-Tspan15 contact.

The most striking feature of the structure is the conformational reorganization of ADAM10 upon complex formation with Tspan15. In the crystal structure of the isolated ADAM10 ectodomain, the protein is in a closed, autoinhibited conformation in which the metalloproteinase domain directly contacts the cysteine-rich domain (Seegar et al., 2017) (Figure 1C, cartoon representation), whereas in the structure of the ADAM10-Tspan15 complex, ADAM10 is held in an open, active conformation by the large extracellular loop (LEL) of Tspan15, which interposes itself between the cysteine-rich and metalloproteinase domains (Figure 1B; see supplementary video S1 for a morph movie comparing the two conformations of ADAM10).

Tspan15, on the other hand, adopts a closed conformation in the complex, with its LEL sitting over the opening created by its four TM helices. This architecture resembles that of tetraspanins CD9 (Umeda et al., 2020), CD53 (Yang et al., 2020) and CD81 (Zimmerman et al., 2016) in isolation (Figure S3, related to Figure 1), and differs from that of CD81 in complex with CD19, the only other bound-state structure of a tetraspanin: in the CD81-CD19 complex, the LEL undergoes a hinge movement to open up by 60° relative to the membrane plane, and the TM helices move closer together to occlude the intramembrane cavity (Susa et al., 2021).

### Structure-function studies of the complex

The LEL domain of Tspan15 is wedged between the metalloproteinase and cysteine-rich domains of ADAM10, forming two discontinuous contact interfaces. The first, here called site A, is between Tspan15 and the ADAM10 cysteine-rich domain, and the second, site B, is between Tspan15 and the ADAM10 catalytic domain (Figure 2A, see Figure S4, related to Figure 2 for electron density of interface regions of each protein, and for the electron density of the bound active-site inhibitor BB94). Previous work has established that a complex still assembles in the absence of the catalytic domain, showing that the site B interface is not required for formation of the complex (Noy et al., 2016b). We therefore focused on mutationally interrogating site A to identify key contacts required for ADAM10 binding to Tspan15.

**Figure 2.**
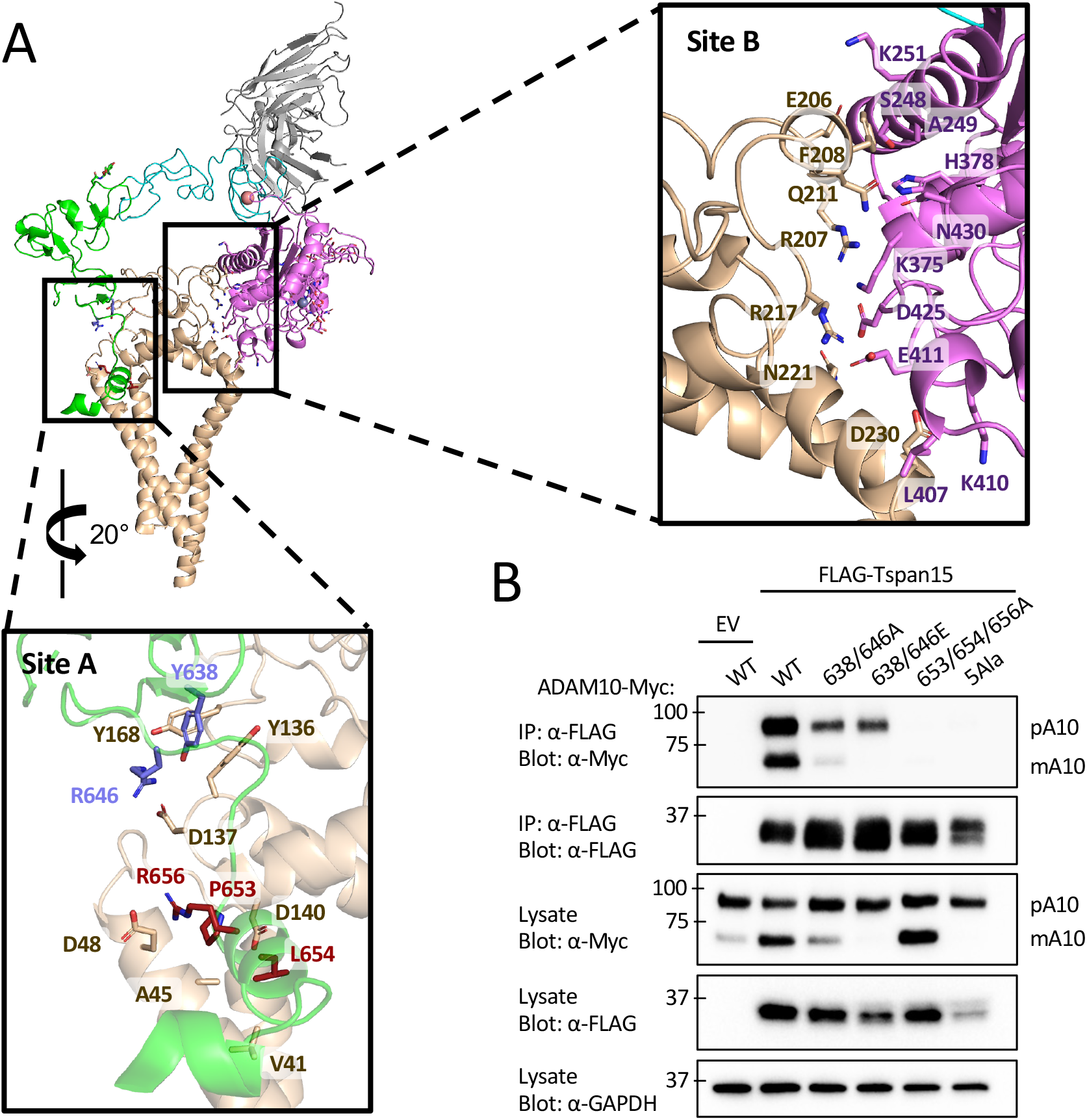
ADAM10-Tspan15 contact interfaces and mutational analysis. (A) Cartoon representation of the vFab-ADAM10-Tspan15 complex with site A (left) and site B (right) interfaces boxed. A close-up view of the interface at site A is shown below the main panel, and a close-up view of the site B interface is shown to the right. The two clusters of interface A residues mutated in the immunoprecipitation assays are shown in blue and maroon. (B) Effect of ADAM10 mutations on co-immunoprecipitation with Tspan15. HEK293 cells were co-transfected with wild-type or mutant ADAM10-myc and FLAG-Tspan15 or vector control, and cell lysates were immunoprecipitated with anti-FLAG antibodies. The lysates and immunoprecipitates were subjected to SDS-PAGE, and analyzed by western blot using anti-myc or anti-FLAG antibodies. Lysates were also probed with an anti-GAPDH antibody (bottom) as a loading control. See also Figure S4.

We introduced a series of mutations into ADAM10 at two patches in the site A interface and evaluated the effects of those changes on ADAM10-Tspan15 complex formation in a coimmunoprecipitation assay. The first patch, mutated at residues Y638/R646, alters a residue pair in contact with Tspan15 residues identified by mutation in a prior study to contribute to ADAM10 binding (Lipper et al., 2022). The second mutation patch, P653/L654/R656, is in a short helix at the membrane surface that contacts the LEL, SEL and TM regions of Tspan15. We made two different sets of mutations in the first patch (Y638A/R646A and Y638E/R646E) and a triple alanine mutation in the second patch (P653A/L654A/R656A). Additionally, we made a five-Ala mutant combining the mutations in both patches (Y638A/R646A/P653A/L654A/R656A). Plasmids expressing Myc-tagged ADAM10 and FLAG-Tspan15 or empty vector were co-transfected into HEK293T cells, and cell lysates were subjected to co-immunoprecipitation as described previously (Lipper et al., 2022) (Figure 2B). WT ADAM10 robustly immunoprecipitated with Tspan15 and showed increased conversion to the mature, processed form when co-transfected with Tspan15 compared to empty vector. Both Y638A/R646A and Y638E/R646E reduced the amount of ADAM that immunoprecipitated with Tspan15, and either reduced (Y638A/R646A) or prevented (Y638E/R646E) processing of ADAM10 into its mature form, respectively. The P653A/L654A/R656A protein failed to detectably co-immunoprecipitate with Tspan15, but retained the ability to undergo processing to the mature form. This observation suggests that this ADAM10 mutant is capable of an interaction (either a weak, transient association with Tspan15 or with another TspanC8) that supports its maturation into the mature form. The 5 Ala mutant (Y638A/R646A/P653A/L654A/R656A) neither immunoprecipitated with Tspan15 nor underwent processing to the mature form, indicating that it was both completely deficient in binding to Tspan15 and incapable of supporting ADAM10 maturation. Together, these data show that Tspan15 binding and efficient ADAM10 maturation rely on these contacts at the site A interface.

### Constraints dictating membrane-proximal cleavage site selectivity

Although the site B interface does not control association of ADAM10 with Tspan15, it positions the active site Zinc ion ~20 Å from the membrane surface (Figure 3A). The active site cleft, occupied in our structure by the hydroxamic acid inhibitor BB94, is also oriented so that the C-terminal end of a bound substrate would be pointed toward the plasma membrane. These two structural features of the complex are consistent with the idea that Tspan15 binding constrains the position and orientation of the ADAM10 active site. For an unstructured 10-residue peptide, the radius of gyration would be approximately 20 Å (R_g_ = nl^2^/6, where n=10 and the estimated Cα-Cα distance l ~ 3.8 Å); therefore, a 20 Å length constraint imposed by Tspan15 would result in preference for a cleavage position about 10 residues from the membrane, assuming that the segment between the membrane and the active site is natively disordered.

**Figure 3.**
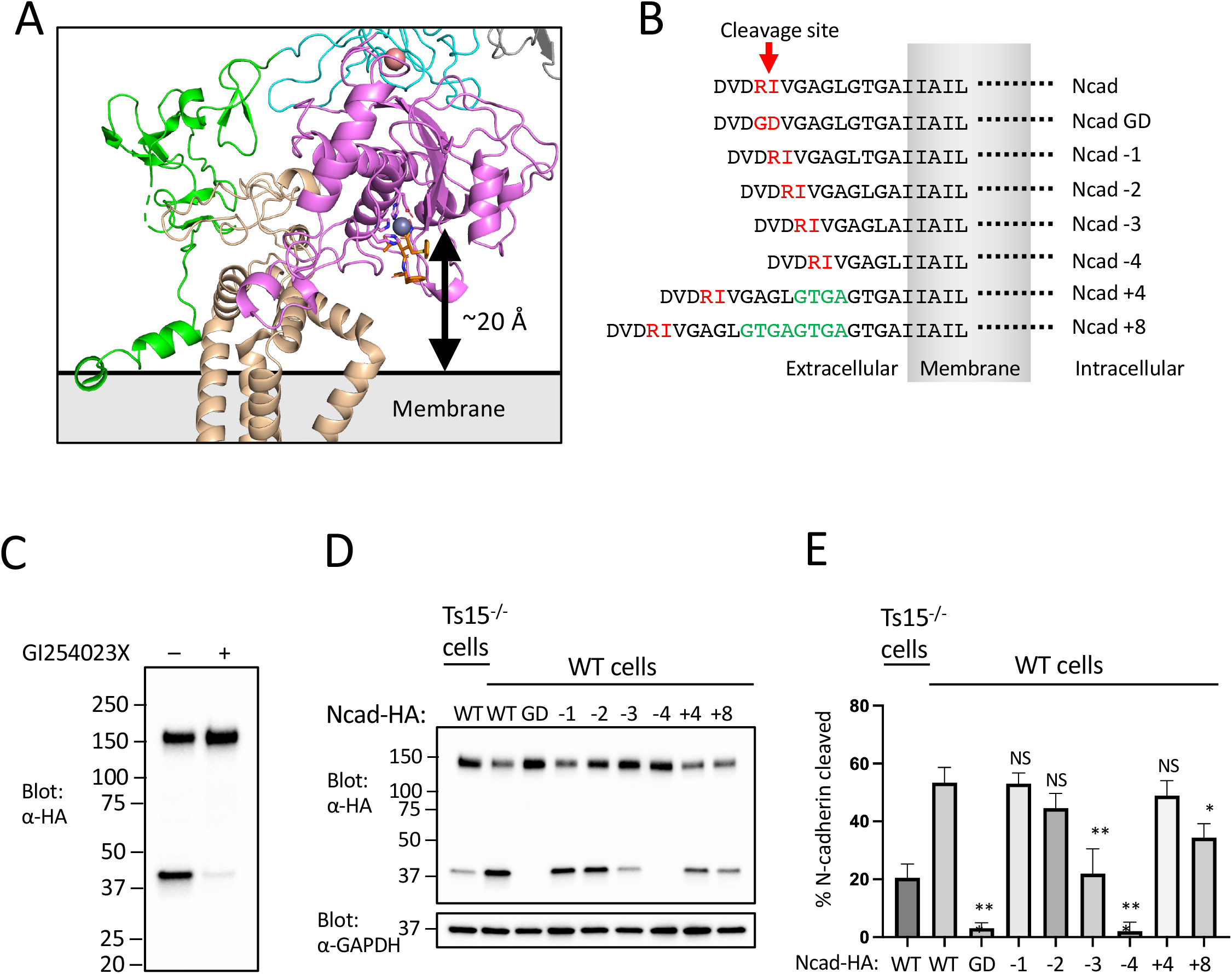
Analysis of the dependence of substrate cleavage on the distance of the scissile bond from the membrane. (A) Zoomed-in view of the ADAM10 active site and its position relative to the membrane. The active site residues are shown in magenta sticks, the catalytic Zn^++^ ion is a gray sphere, and the BB94 hydroxamic acid inhibitor is rendered in orange sticks. (B) Design of N-cadherin (Ncad) variants for the substrate cleavage assay, and nomenclature used to name the substrate variants. (C) ADAM10 dependence of N-cadherin cleavage. U251 cells were transfected with Ncad containing an HA tag it its C-terminal end, and either mock treated or treated with the ADAM10 inhibitor GI254023X. Cell lysates were subjected to SDS-PAGE, and analyzed by Western blot using an anti-HA antibody. (D) N-cadherin substrate cleavage assay. Parental or Tspan15 knockout (Ts15^-/-^) U251 cells were transfected with one of the HA-tagged Ncad variants shown in panel B. Cells were lysed after 48 hours and blotted with an anti-HA antibody to determine the extent of cleavage. Lysates were also blotted with anti-GAPDH (bottom) as a loading control. Data are representative of three independent biological replicates. (E) Quantification of N-cadherin cleavage using the Western blot results from all replicates. Error bars indicate standard deviation. * p< 0.05; ** p<0.01; *** p<0.001.

To test this model, we varied the distance between the ADAM10 cleavage site and the membrane for the substrate N-cadherin (Ncad) and monitored the effects of changing the position of the processing site on the efficiency of cleavage by ADAM10. The wild-type (WT) Ncad cleavage site is 10 residues from the plasma membrane. We designed a series of deletions and insertions in the membrane proximal region of Ncad to create variants with cleavage sites ranging from 6-18 residues from the membrane, along with a cleavage-resistant control mutant Ncad-GD (Uemura et al., 2006) (Figure 3B). We analyzed the extent of Ncad cleavage of the different length variants in U251 cells, which rely on ADAM10 (Figure 3C) and Tspan15 for this processing event (Lipper et al., 2022) (Figure 3D,E). The proteolysis results show that cleavage efficiency is optimal when the processing site is between eight and fourteen residues from the membrane. When the separation between the processing site and the membrane was shorter than nine residues, cleavage efficiency decreased, and when the distance was shortened to six residues (Ncad-4) processing was indistinguishable from the Ncad-GD cleavage resistant mutant (Figure 3D,E). As predicted from a random walk model, increasing the distance between the processing site and the membrane was better tolerated, with a four-residue insertion (Ncad+4) exerting no significant effect and only the eight-residue insertion (Ncad+8) showing a small, but significant, reduction in cleavage efficiency (Figure 3D,E).

Next, we tested whether site B is required for selective cleavage of the WT Ncad substrate by testing the effect of mutating the site B interface directly (Figure 4A). Disruption of the site B interface with a trio of point mutations further disfavors cleavage of shorter substrates without affecting cleavage of long substrates (Figure 4B,C), as predicted by a model in which site B tethers the ADAM10 active site proximal to the membrane.

**Figure 4.**
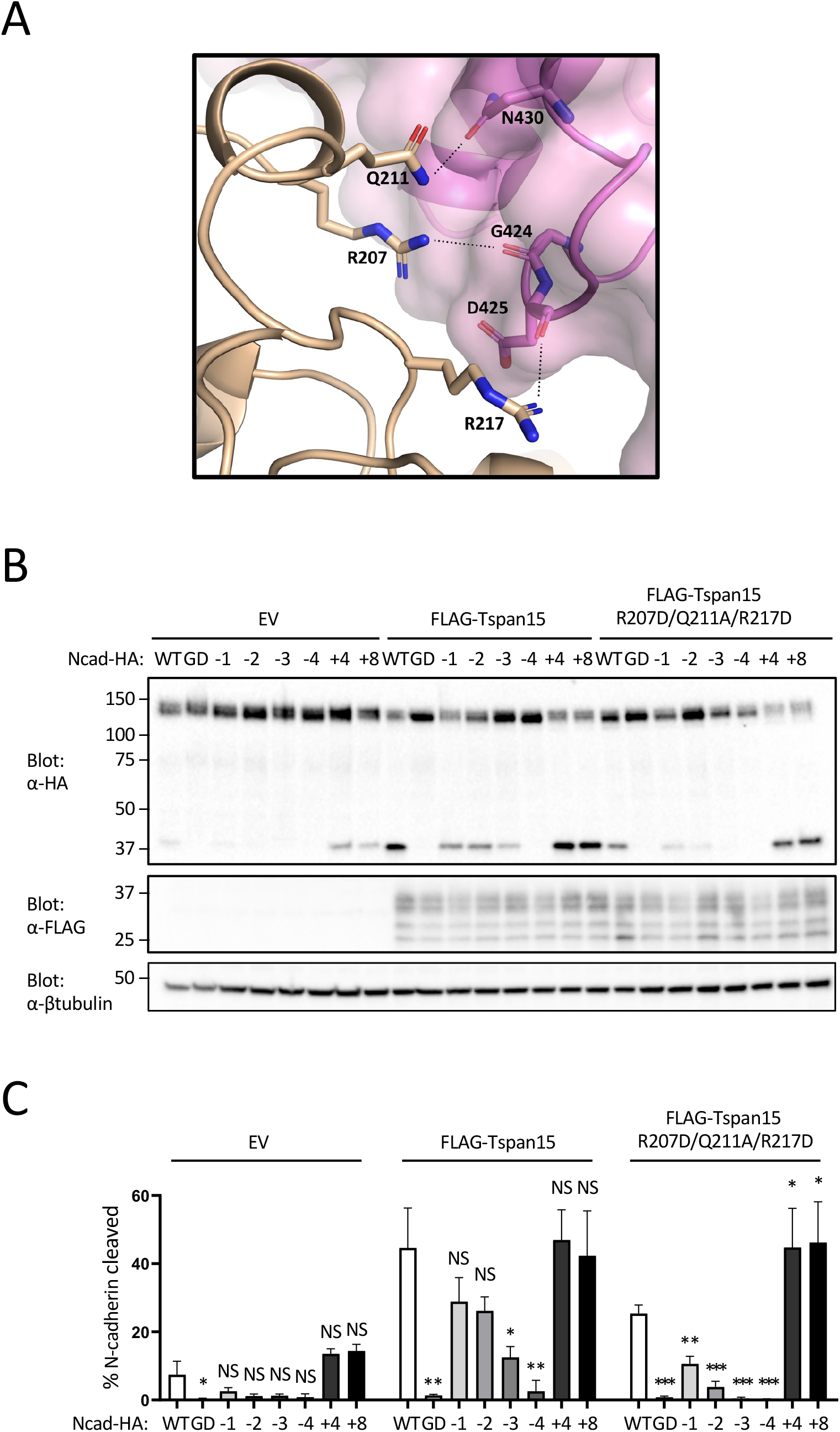
Analysis of the Tspan15-catalytic domain (site B) interface. (A) View of the ADAM10-Tspan15 complex highlighting the residues mutated at the site B interface. Tspan15 is shown in beige, and that catalytic domain of ADAM10 is shown in magenta. Side chains of Tspan15 interface residues that were mutated (R207D, Q211A, and R217D) are shown as sticks. ADAM10 residues shown as sticks are within H-bonding distance (dashed lines) of Tspan15. (B) N-cadherin cleavage assay analyzing the Tspan15 site B interface mutant. Tspan15 knockout U251 cells were transfected with vector control (EV), FLAG-tagged wild-type Tspan15 or the Tspan15 site B interface mutant, and with one of the HA-tagged Ncad variants. Cells were lysed after 48 hours, and blotted with anti-HA to determine the extent of cleavage, and with anti-FLAG to confirm Tspan15 protein expression. Lysates were also blotted with anti-β-tubulin (bottom) as a loading control. Data are representative of three independent biological replicates. (C) Quantification of N-cadherin cleavage using the Western blot results from all replicates. Error bars indicate standard deviation. * p< 0.05; ** p<0.01; *** p<0.001. See also Figure S5.

The residues on Tspan15 at the contact interface with the metalloproteinase domain of ADAM10 vary among the six TspanC8 partner proteins that direct differential cleavage selectivity of ADAM10 for substrates (Figure 5A). In particular, Tspan5 and Tspan14 are divergent from Tspan15 in this region, have been shown to favor cleavage of Notch substrates, and disfavor cleavage of Ncad, suggesting that differences in this region among these TspanC8 proteins might alter positional selectivity of ADAM10 for substrate cleavage sites. To test this possibility, we designed chimeras of Tspan15 with Tspan5 and with Tspan14 that swap the sequence at the metalloprotease interface, spanning residues 206-230. The effect of these sequence swaps on N-cadherin cleavage was then analyzed using WT Ncad and the elongated Ncad+4 as substrates. Unlike Tspan15, which promoted similar levels of cleavage for both WT Ncad and Ncad+4, the Tspan15-Tspan5 and Tspan15-Tspan14 chimeras exhibited impaired cleavage of Ncad while promoting cleavage of Ncad+4 more efficiently than the parental Tspan15 protein (Figure 5B,C). Consistent with our model, these results reveal that selectivity in substrate processing by distinct ADAM10-TspanC8 complexes can be tuned by the distance of the substrate cleavage site from the membrane.

**Figure 5.**
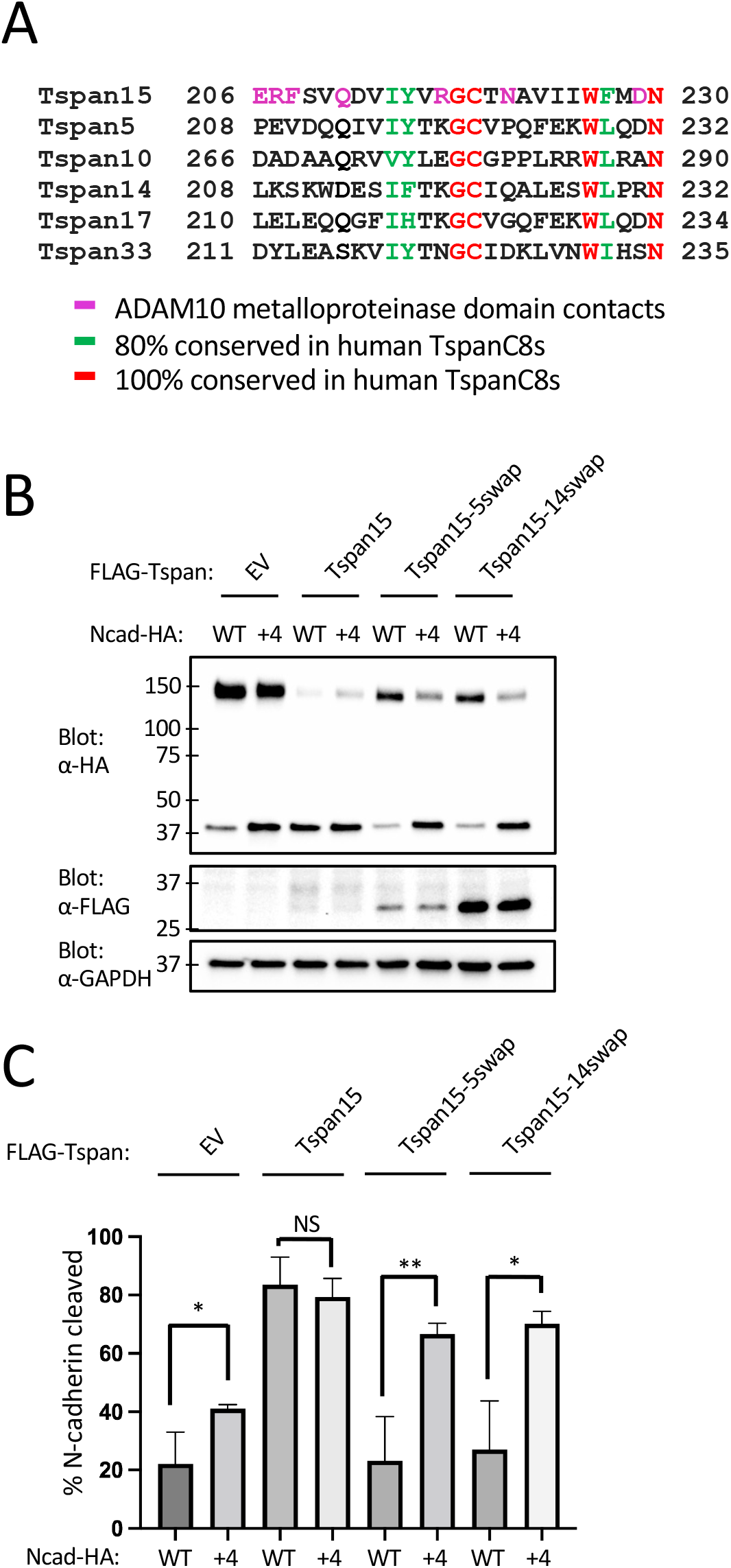
N-cadherin cleavage assay analyzing Tspan site B chimeras. (A) Sequence alignment of the site B interface region of the six C8-tetraspanin proteins. (B). Tspan15 knockout U251 cells were transfected with vector control (EV), FLAG-tagged wild-type Tspan15, Tspan15/Tspan5, or Tspan15/Tspan14 chimeric proteins, and with either an HA-tagged Ncad or Ncad+4 variant. Cells were lysed after 48 hours, and blotted with anti-HA to determine the extent of cleavage, and with anti-FLAG to confirm Tspan protein expression. Lysates were also blotted with anti-GAPDH (bottom) as a loading control. Data are representative of three independent biological replicates. (C) Quantification of N-cadherin cleavage using the Western blot results from all replicates. Error bars indicate standard deviation. * p< 0.05; ** p<0.01. See also Figure S5.

## Discussion

The structural and cell-based work reported here uncover the basis for intrinsic and extrinsic mechanisms that influence substrate selectivity of the alpha secretase ADAM10. In the structure of the isolated ectodomain, ADAM10 adopts a partially autoinhibited conformation and the active site of the catalytic domain has a deep hydrophobic pocket selective for bulky hydrophobic residues at the P1’ position of the substrate (Seegar et al., 2017). In our cryo-EM structure of the Tspan15-ADAM10 complex, the bound Tspan15 converts ADAM10 into an open conformation and situates the catalytic site of the enzyme ~20 Å from the membrane surface, effectively creating a molecular ruler for cleavage at membrane proximal substrate sites.

The positioning of the ADAM10 catalytic site relative to the membrane is defined by the presence of two ADAM10-Tspan15 interfaces at site A and site B. Whereas the sequence at the site A interface is conserved among the C8 tetraspanins, the sequence at the site B interface is not. Emerging data suggest that complexation of ADAM10 with the six different C8-Tspan proteins creates six distinct molecular species with differential cleavage selectivity for substrates such as VE- and N-cadherins, Notch receptors, and APP. Among well characterized ADAM10 substrates, the ADAM10-Tspan15 complex selectively cleaves Ncad (Noy et al., 2016b) and platelet glycoprotein (GP) VI (Koo et al., 2022), and Tspan5 and Tspan14 promote Notch processing by ADAM10 (Eschenbrenner et al., 2020; Jouannet et al., 2016). In Ncad and GPVI, the P1’ cleavage site is 10 residues from the membrane (note that misassignment of the start of the transmembrane region of GPVI by the Uniprot database has misled others (Koo et al., 2022) into stating that its cut site is five residues from the membrane, rather than 10), optimized for processing by ADAM10-Tspan15, whereas the separation of the cut site from the membrane is 15 residues for human Notch1 (Mumm et al., 2000). While it is possible that different patterns of localization or direct substrate recruitment by different TspanC8 proteins may account for some differences in the cleavage preferences of various ADAM10-TspanC8 complexes, our structural and proteolytic data suggest that differences in cleavage patterns among the six ADAM10-TspanC8 complexes also result from differences in (or the absence of) the site B interface, which is not conserved among the TspanC8 proteins.

Modulation of ADAM10 activity has therapeutic potential for a number of human diseases ranging from Alzheimer’s disease to cancer (Liu et al., 2006; Loganathan et al., 2020; Sulis et al., 2011; Venkatesh et al., 2017; Weskamp et al., 2006). The development of therapeutics that act directly on ADAM10 has been challenging due to its numerous physiological substrates and the subsequent on-pathway toxicity that results from altering their processing (Wetzel et al., 2017). Our findings open up the possibility of selectively inhibiting or activating cleavage of a distinct target protein such as Notch1 or APP using an antibody directed at a specific TspanC8 protein, analogous to the modulation of CD19-CD81 complexation in B cell co-receptor signaling with an antibody directed at the Tspan CD81 (Susa et al., 2020). Promisingly, the anti-Tspan15 antibody 1C12 was able to bind Tspan15 when in complex with ADAM10 and displayed a partial inhibitory effect on ADAM10 cleavage of VE-cadherin in a cell-based assay (Koo et al., 2020). When superimposed on the ADAM10-Tspan15 complex structure reported here, our structure of the 1C12 Fab bound to the Tspan15 LEL (Lipper et al., 2022) shows that the Fab would clash with the disintegrin and metalloproteinase domains (Supplementary Figure S5, related to Figures 3–5), indicating that the antibody likely displaces the catalytic domain of ADAM10 from Tspan15 to interfere with VE-cadherin cleavage. Development of TspanC8-directed antibodies targeting the site B interface could be developed to modulate ADAM0 substrate selectivity for therapeutic applications, such as to promote alpha-secretase cleavage of APP. Other antibodies that promote or inhibit the activity of distinct ADAM10-TspanC8 complexes should enable specificity in modulating ADAM10 activity toward other desired substrates (*e.g*. Notch, N-cadherin) while reducing the undesired consequences of modulatory antibodies or compounds that directly activate or inhibit ADAM10.

### Limitations of the Study

One caveat to note is that our structure was determined in detergent, not in the presence of membrane lipids or in an intact bilayer. Although we did show that the complex is active in cleaving a fluorogenic peptide substrate in the identical buffer to that used for structure determination, our studies do not provide any information about potential structural roles for lipids in complex formation or stabilization of either the primary (site A) or secondary (site B) interface. In addition, the transmembrane portion of ADAM10 was also not visible in the structure, and it is possible that this region becomes ordered in the natural microenvironment of the membrane. Another limitation we encountered was that it was not possible to test the Tspan15 length-dependence for cleavage of APP and Notch substrates in our U251 cell-based assay system because there was a high amount of basal cleavage activity toward those substrates even in Tspan15 knockout cells. Nevertheless, our studies provide a clear structural rationale for the length constraint in selective cleavage of membrane-proximal substrates by ADAM10-Tspan15 complexes.

## Acknowledgments

We thank the cryo-EM Center for Structural Biology at Harvard Medical School for expert assistance with cryo-EM data acquisition and processing, and members of the Blacklow lab for helpful discussions. Financial support for this work was provided by NCI grants R35 CA220340 and R01 AI172846 (to S.C.B.), and a gift from Edward B. Goodnow (to S.C.B.).

## Author Contributions

Experimental Design: C.H.L., E.D.E., and S.C.B. Materials preparation: K-H.G., C.H.L. and E.D.E. Data acquisition: C.H.L. and E.D.E. Data analysis and interpretation: C.H.L. E.D.E., and S.C.B. Manuscript preparation and editing: C.H.L. and S.C.B. with input from E.D.E. Acquisition of Funding: S.C.B.

## Inclusion and Diversity

One or more of the authors of this paper self-identifies as an underrepresented ethnic minority in their field of research or within their geographical location.

## Supplementary Figure Legends

**Figure S1.**
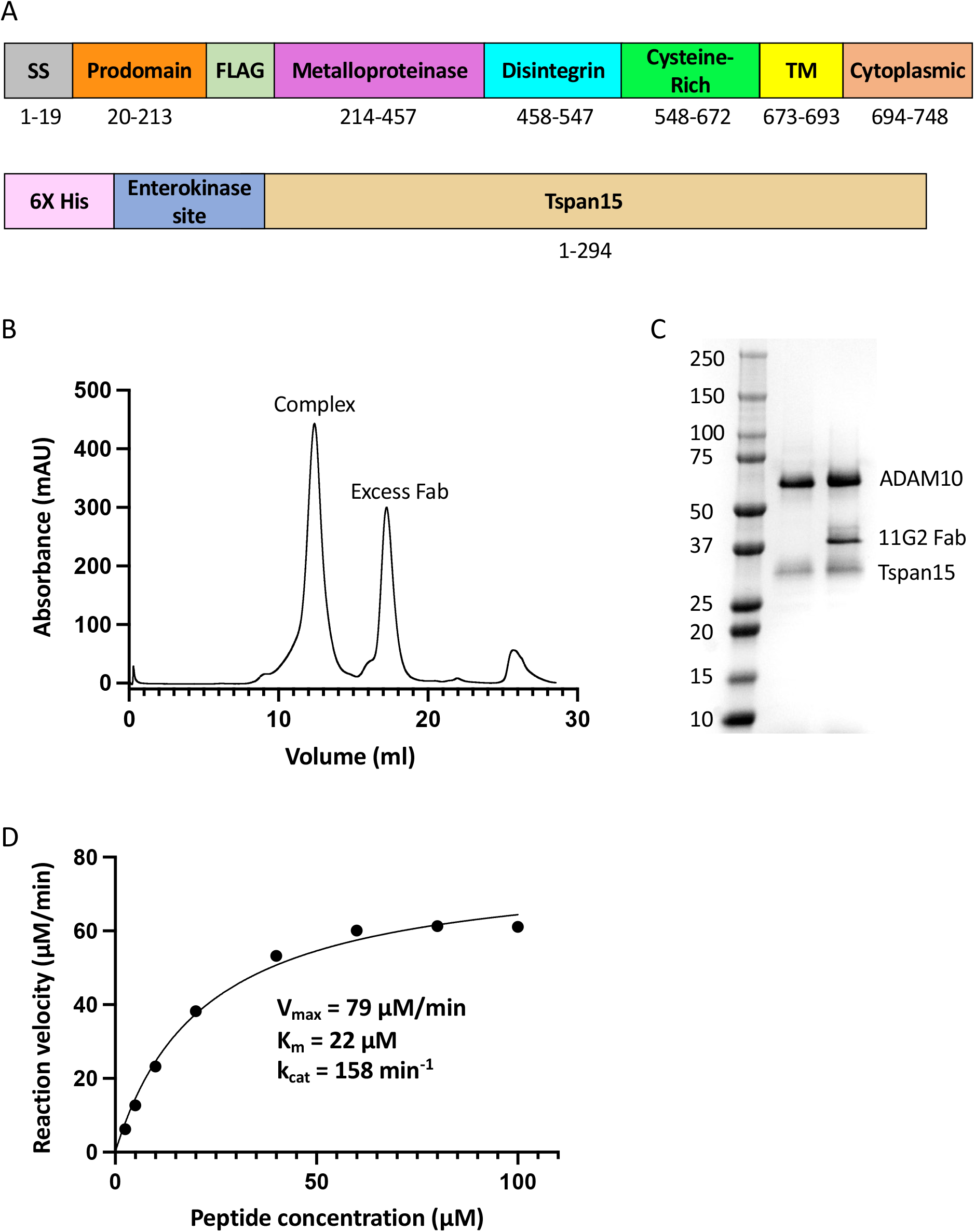
Construct design, protein purification, and enzymatic activity, related to Figure 1. (A) Domain organization of the ADAM10 and Tspan15 proteins produced for structural studies. Note the placement of a FLAG tag inserted between the prodomain and the metalloproteinase domain of ADAM10 to facilitate purification of the mature species, from which the prodomain has been released. (B) Size exclusion chromatogram of the purified 11G2 Fab-ADAM10-Tspan15 complex, isolated on a S200 column. (C) SDS-PAGE analysis of purified ADAM10-Tspan15 and purified 11G2 Fab-ADAM10-Tspan15 complexes, visualized by staining with Coomassie blue dye. (D) Enzymatic activity of the purified ADAM10-Tspan15 complex, using the fluorogenic peptide substrate, Mca-PLAQAV-Dpa (R & D systems).

**Figure S2.**
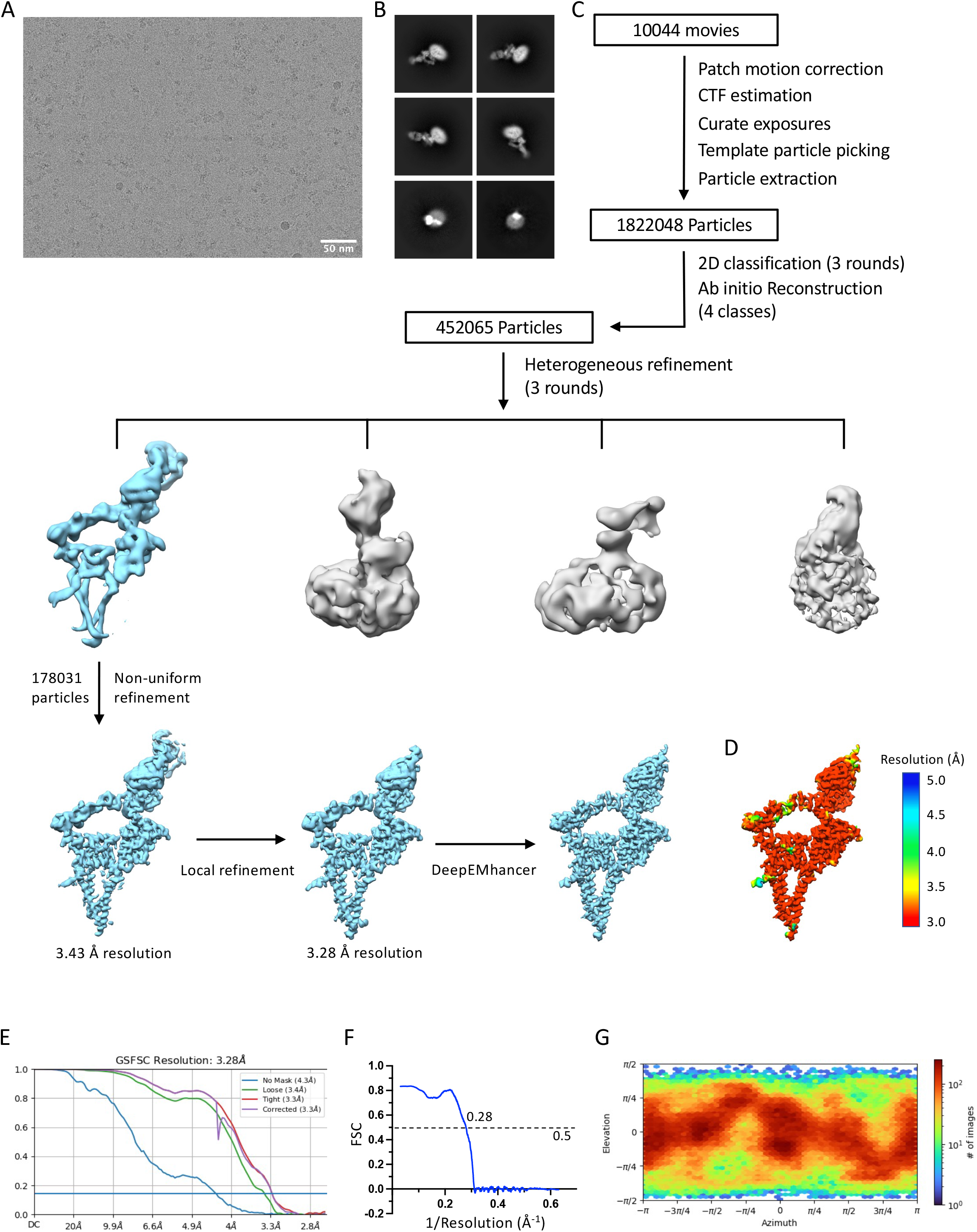
Cryo-EM image processing workflow, related to Figure 1. (A) A representative raw cryo-EM image. Particles were extracted with a box size of 300 pixels and pixel size of 1.32 Å. (B) Two-dimensional class averages of the Fab-ADAM10 Tspan15 complex. (C) Cryo-EM data processing workflow. (D) Density map colored by local resolution according to the scale at the right of the panel. (E) Fourier shell correlation (FSC) used to determine overall resolution of the map. The blue line marks the resolution corresponding to an FSC value of 0.143. (F) Map to model FSC curve prepared using Phenix Comprehensive Validation tool. The value where the curve crosses the dotted line at 0.5 is labeled. (G) Angular distribution of the Fab-ADAM10 Tspan15 particles in the final round of 3D refinement in CryoSPARC.

**Figure S3.**
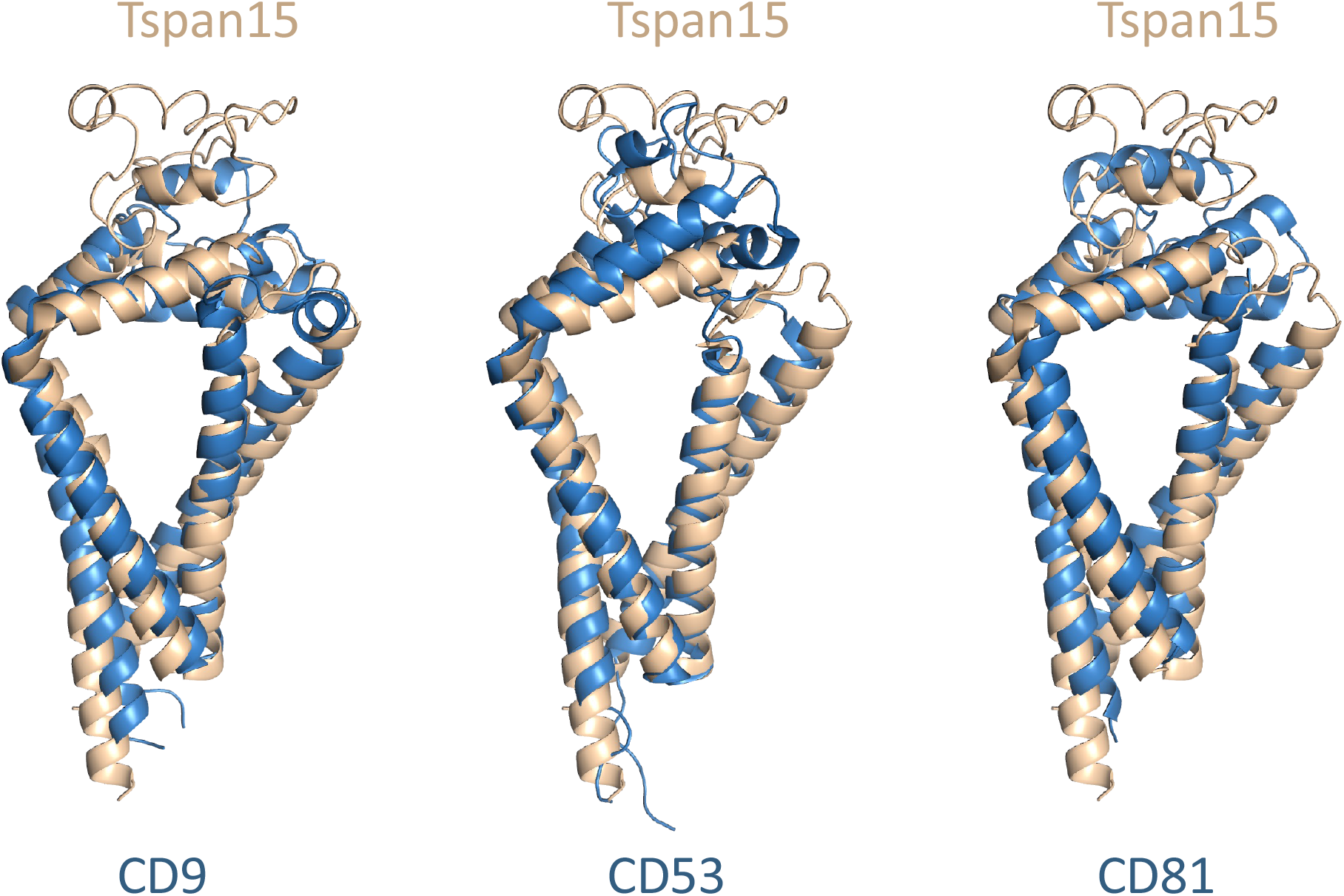
Structural comparisons of Tspan15 in the complex with the CD9 (left), CD53 (center), and CD81 (right) in their closed conformations. Tspan is in beige, and the other tetraspanins are in blue. Related to Figure 1.

**Figure S4.**
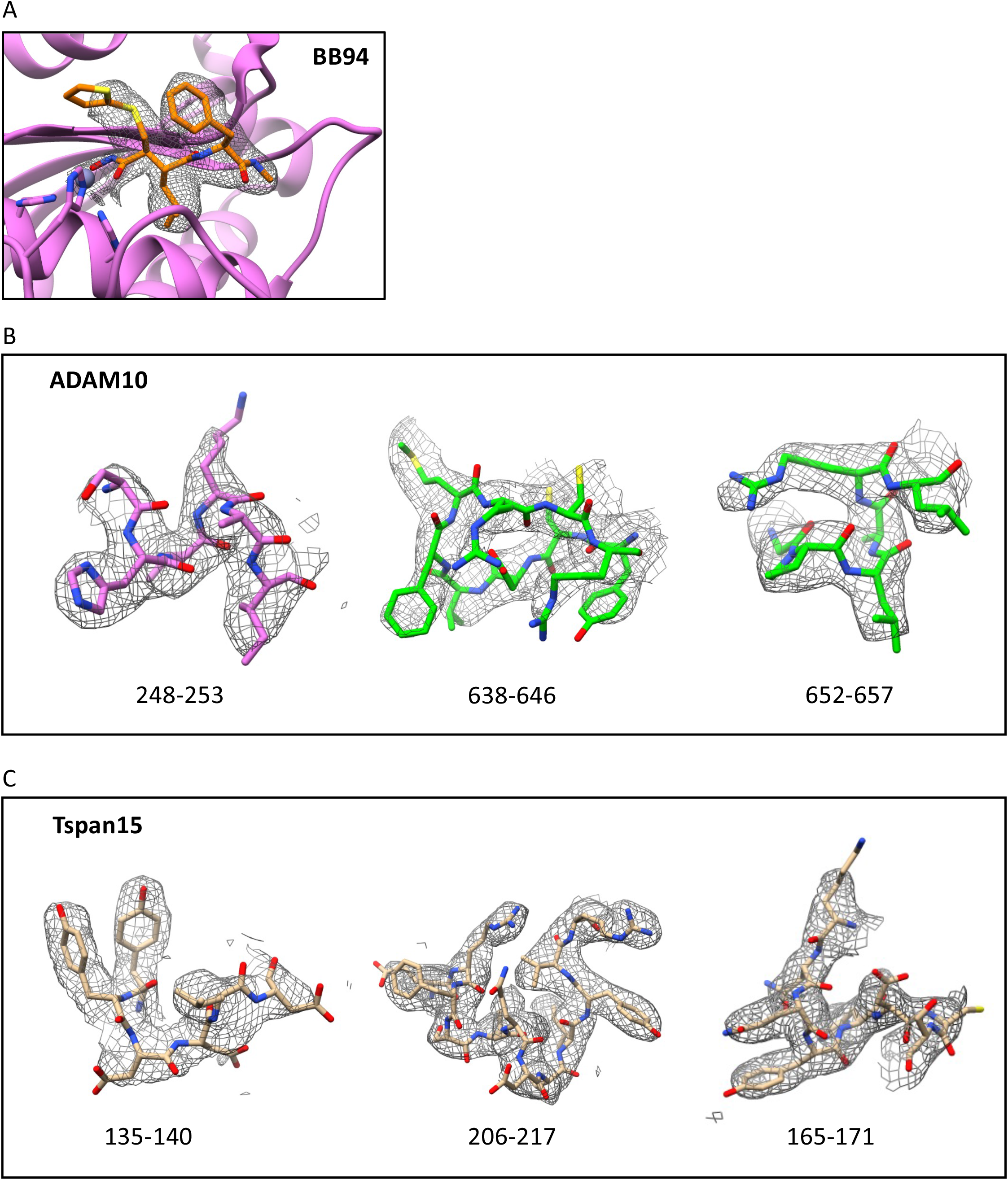
Representative electron density at various sites in the complex, related to Figure 2. (A) Electron density around the hydroxamic acid inhibitor BB94 (gray mesh). The inhibitor is rendered as sticks in orange and CPK colors. The catalytic domain of ADAM10 is shown as a ribbon diagram in magenta. (B). Electron density in three regions of ADAM10 that form contacts with Tspan15. Left panel: density around residues 248—253 in the catalytic domain. The density is shown in gray mesh and the polypeptide segment is shown as sticks in magenta and CPK colors. Center and right panels: density around residues 638—646 and 652-657, respectively, in the cysteine-rich domain near the contact site. The density is shown in gray mesh and the polypeptide segment is shown as sticks in green and CPK colors. (B). Electron density in three regions of Tspan15 that form contacts with ADAM10. Left, center and right panels: density around residues 135—140, 206-217 and 165-171. The density is shown in gray mesh and the polypeptide segment is shown as sticks in beige and CPK colors.

**Figure S5.**
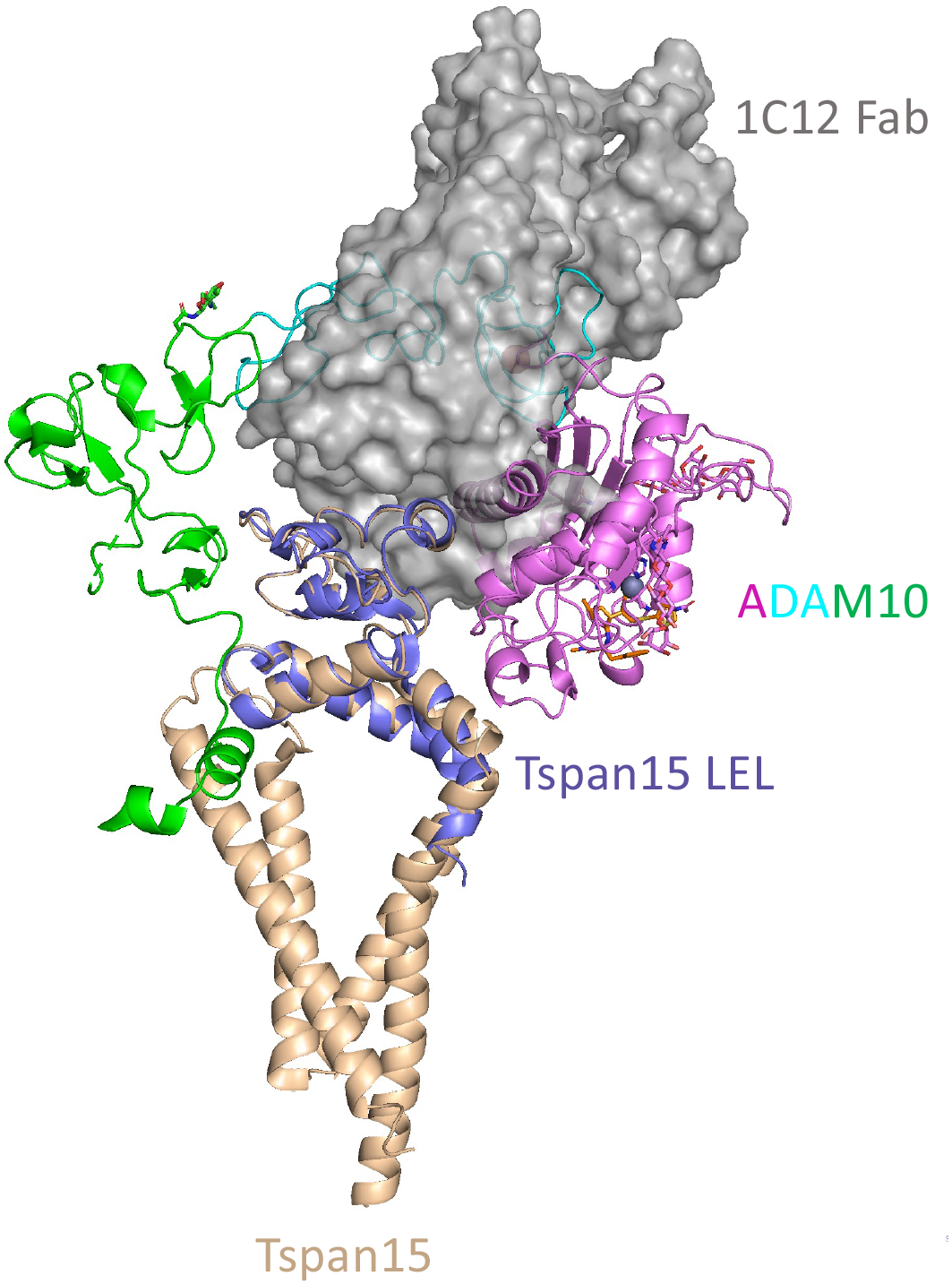
Predicted steric clash between the Tspan15-bound 1C12 Fab and ADAM10 in the Tspan15-ADAM10 complex, related to Figures 3–5. The structure of the Tspan15-ADAM10 complex is rendered as ribbons according to the color scheme in Figure 1. This model is superimposed on the structure of the Tspan15 ectodomain (purple ribbon representation) in its complex with the 1C12 Fab, which is shown as a gray surface. Note the steric clashes between i) the Fab and the catalytic domain (purple) and ii) the Fab and the disintegrin domain (blue).

**Table S1.** Data Collection and Refinement Statistics, related to Figure 1.

|  |  |
| --- | --- |
| **Data collection and processing** |  |
| Magnification | 105,000 |
| Voltage (kV) | 300 |
| Total dose (e-/Å<sup>2</sup>) | 51.99 |
| Dose per physical pixel per second | 27.22 |
| Dose per Å<sup>2</sup> per second | 39.993 |
| Exposure time (s) | 1.3 |
| Number of frames | 50 |
| Dose per frame (e-/Å<sup>2</sup>) | 1.04 |
| Defocus range (μm) | -1.0 to -2.2 |
| Pixel size (Å) | 0.825 |
| Symmetry | C1 |
| Initial particle images | 1822048 |
| Final particle images | 178031 |
| Map resolution (Å) | 3.3 |
| FSC Threshold | (0.143) |
| **Refinement** |  |
| Model composition |  |
| Non-hydrogen atoms | 7222 |
| Protein residues | 933 |
| Ligands | 13 |
| B-factors (Å<sup>2</sup>) |  |
| Protein | 56.68 |
| Ligands | 68.31 |
| R.m.s deviations |  |
| Bond lengths (Å) | .003 |
| Bond angles (°) | 1.036 |
| Validation |  |
| MolProbity score | 2.06 |
| Clashscore | 12.85 |
| Rotamer outliers (%) | 0 |
| Ramachandran plot |  |
| Favored (%) | 93.07 |
| Allowed (%) | 6.72 |
| Disallowed (%) | 0.22 |

## STAR Methods

### Key resources table

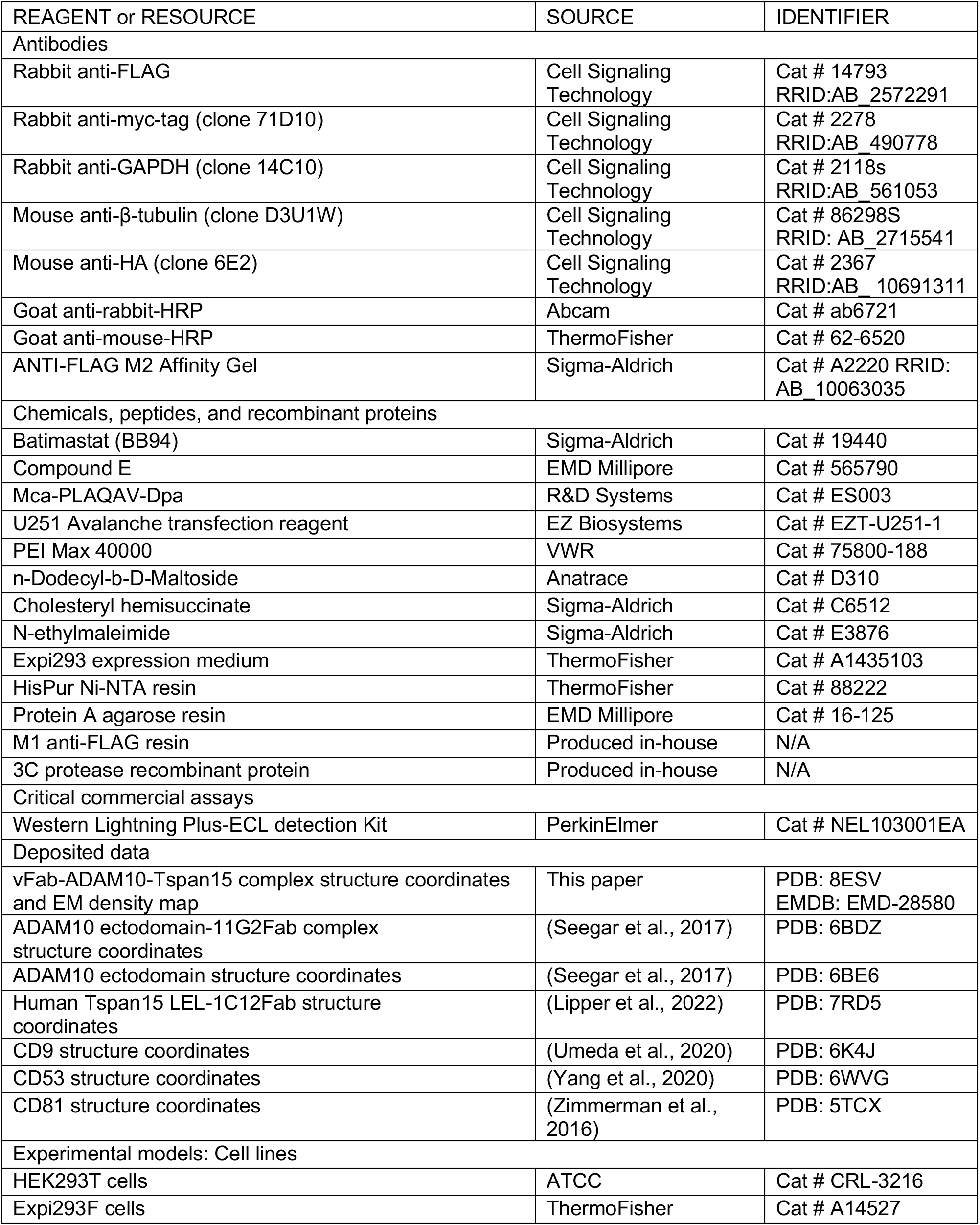
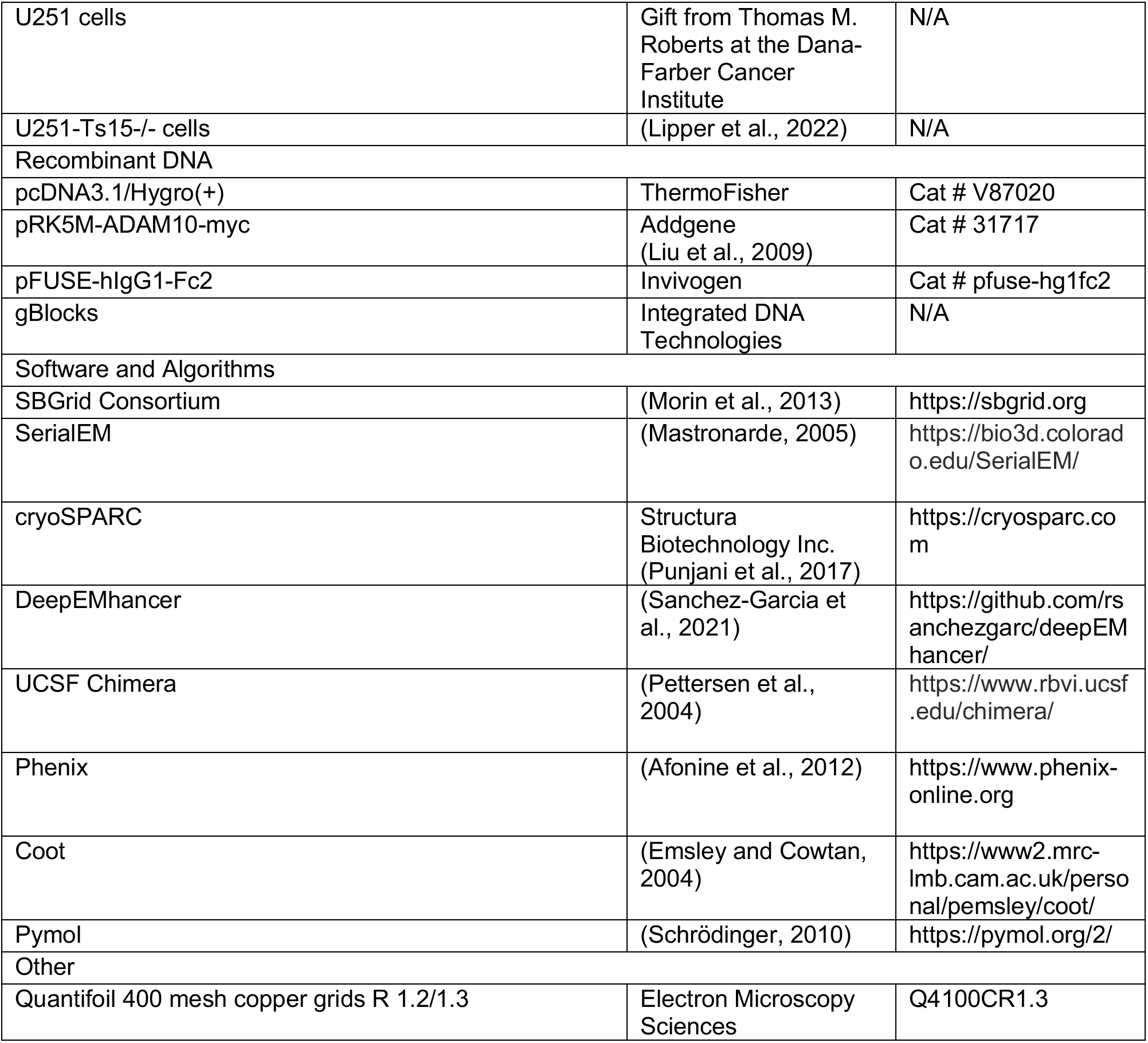

### Resource availability

#### Lead contact

Further information and requests for either resources or reagents should be directed to and will be fulfilled by the lead contact Stephen C. Blacklow.

#### Materials availability

Requests for materials generated in this study should be directed to and will be fulfilled by the lead contact Stephen C. Blacklow.

#### Data and Code Availability

The coordinates of the 11G2 vFab-ADAM10-Tspan15 complex have PDB ID code 8ESV and the EM map has EMDB accession number EMD-28580. Other materials are available upon request to S.C.B. This paper does not report original code. Any additional information required to reanalyze the data reported in this paper is available from the lead contact upon request.

### Experimental model and subject details

Protein for cryo-EM was isolated from Expi293F cells. Expi293 cells were maintained in suspension at 37°C, 8% CO_2_ in Expi293 expression medium (ThermoFisher). Co-immunoprecipitation experiments were performed in HEK293T cells. N-cadherin cleavage assays were performed in U251 cells. HEK293T and U251 cells were maintained at 37°C in DMEM (Corning) supplemented with 10% fetal bovine serum (GeminiBio) and penicillin/streptomycin (Gibco).

### Method details

#### Plasmid construction

ADAM10 used for structure determination was expressed using a modified version of the PRK5M-ADAM10 plasmid (Addgene plasmid 31717) with a FLAG tag immediately following the prodomain boundary site and the myc tag removed (FLAG-ADAM10). Tspan15 was cloned into pcDNA3.1/Hygro(+) with an N-terminal 6-histidine tag followed by an enterokinase cleavage site (used as a linker, His-EK-Tspan15). A single DNA insert containing the 11G2 light chain followed by a P2A ribosomal skipping sequence and then the heavy chain Fab sequence was inserted into the pFUSE-hIgG1-Fc2 plasmid containing the human IgG Fc. The sequence contains a 3C protease site between the light chain and the P2A sequence, and another 3C site between the heavy chain and the Fc. The light chain uses the IL2 signal sequence in the vector and the heavy chain uses the native signal sequence. 11G2 contains the N79Q mutation in the heavy chain to remove a glycosylation site. N-cadherin wild-type and mutant sequences were cloned into pcDNA3.1/Hygro(+) with a C-terminal HA tag. Tspan15-Tspan5 and Tspan15-Tspan14 chimeras replace Tspan15 residues 206-230 with Tspan5 residues 208-232 and Tspan14 residues 208-232, respectively. These constructs as well as wild-type Tspan15 were inserted into pcDNA3.1/Hygro(+) with an N-terminal FLAG tag. Co-immunoprecipitation and N-cadherin cleavage assays used FLAG-Tspan15 constructs and ADAM10-myc constructs (Addgene plasmid 31717 and point mutants derived from it).

#### Protein expression

All proteins were expressed in Expi293F cells. Cells were grown in Expi293 growth medium to a density of 3.0×10^6^ cells/mL and then transfected with 1.0 mg DNA / L of culture using PEI MAX 40K reagent at a 1:3 DNA/PEI ratio. 24 hours after transfection 10 mL per L culture of 45% D-(+)-Glucose solution (Sigma-Aldrich) and 3 mM valproic acid sodium salt (Sigma-Aldrich) were added to the cells. For expression of the ADAM10-Tspan15 complex, cells were co-transfected with each plasmid in a 1:1 ratio and cells were incubated for 48 hours after transfection before harvesting. For 11G2 antibody expression, cells were incubated for 5-6 days after transfection before the antibody was recovered from the media.

#### 11G2 Fab purification

Conditioned media containing 11G2 antibody was passed over Protein A agarose. The retained protein was washed with buffer containing 20 mM HEPES pH 7.4 and 150 mM NaCl. The antibody was eluted with buffer containing 20 mM glycine pH 3.0 and 150 mM NaCl, then neutralized by adding 1 M HEPES, pH 7.4 to a final buffer concentration of 100 mM. The Fc was released from the Fab by adding 3C protease at a ratio of 1:40 (w/w) and incubating at room temperature overnight. The Fc was then removed from the solution by passage over Protein A resin.

#### ADAM10-Tspan15 complex purification

Cells expressing ADAM10 and Tspan15 were harvested by centrifugation and lysed by osmotic shock in 20 mM HEPES pH 7.4, containing 2 mM MgCl2, 10 μM ZnCl2, 1 mM CaCl2, 2 mg/ml iodoacetamide (Sigma Aldrich), 1:50,000 (v:v) benzonase nuclease, and 1 μM BB94. Lysed cells were then centrifuged at 50,000 x g for 20 minutes. The ADAM10-Tspan15 complex was then extracted in buffer containing 20 mM HEPES pH 7.4, 150 mM NaCl, 10 % (v/v) glycerol, 1% (w/v) n-Dodecyl-β-D-Maltoside (DDM), 0.1% (w/v) cholesteryl hemisuccinate (CHS), 2 mg/ml iodoacetamide, 10 mM ZnCl2, 5 mM CaCl2 and 1 μM BB94 using a glass Dounce homogenizer, then stirred for 1 h at 4 °C. Extracted protein was centrifuged at 50,000 x g for 30 minutes. Supernatant was loaded onto a column containing 3 ml M1 FLAG resin, washed with 10 ml of 20 mM HEPES pH 7.4, containing 150 mM NaCl, 1 % glycerol, 0.1% DDM, 0.01% CHS, 5 mM CaCl2, and 1 μM BB94, then washed with 40 ml of 20 mM HEPES pH 7.4, containing 150 mM NaCl, 0.03% GDN, 0.003% CHS, 5 mM CaCl2, and 1 μM BB94. Protein was eluted with 0.2 mg/ml FLAG peptide in 20 mM HEPES pH 7.4, containing 150 mM NaCl, 0.03% GDN, 0.003% CHS, and 1 μM BB94. Eluted protein was then added to Ni-NTA resin and washed with 20 mM HEPES pH 7.4, containing 150 mM NaCl, 0.03% GDN, 0.003% CHS, 20 mM imidazole, and 1 μM BB94. Protein was eluted with the same buffer containing 250 mM imidazole. The ADAM10-Tspan15 complex was incubated with excess 11G2 Fab for 1 h and further purified by size-exclusion chromatography using a Superdex 200 Increase 10/300 GL column in 20 mM HEPES pH 7.4, containing 150 mM NaCl, 0.03% GDN, and 0.003% CHS. ADAM10-Tspan15 used for enzymatic analysis was purified without the addition of BB94 and 11G2 Fab, with final buffer conditions of 20 mM HEPES pH 7.4, 150 mM NaCl, 0.03% GDN, 0.003% CHS, and 2 μM ZnCl2.

#### Fluorogenic peptide cleavage assay

The reaction was initiated by mixing ADAM10-Tspan15 complex (final concentration 0.5 μM) with 25 μM fluorogenic peptide substrate, Mca-PLAQAV-Dpa (R&D Systems) at 37 °C. The reaction buffer consisted of 20 mM HEPES pH 7.4, 150 mM NaCl, 0.03% GDN, 0.003% CHS, and 2 μM ZnCl2. The reaction was monitored by fluorescence emission (excitation = 320 nm and emission = 405 nm) using a SpectraMax M5 Microplate Reader (Molecular Devices). Data were analyzed using Graphpad Prism software. Two replicates were performed.

#### Cryo-EM grid preparation

Quantifoil holey carbon film-coated 400 mesh copper grids (Electron Microscopy Sciences, R1.2/1.3) were glow discharged using a PELCO easiGlow (Ted Pella) for 30 s at 15 mA. A Vitrobot Mark IV was used for sample application and plunge-freezing. 3.3 μl of 2.1 mg/ml ADAM10-Tspan15-11G2 was applied to each grid at 22 °C with 100% humidity, followed by blotting for 7 s with a blot force of 15. The grids were then plunge-frozen in liquid ethane and stored in liquid nitrogen until data collection.

#### Cryo-EM data collection

Grids were imaged on a FEI Titan Krios operated at 300 kV with a K3 summit direct electron detector camera in counting mode. Data were acquired at a nominal magnification of 105,000x, pixel size of 0.825 Å, total exposure dose of 51.99 e^-^/Å^2^ over 50 frames and 1.04 e^-^/Å^2^ dose per frame. A total of 10037 movies were collected with a defocus in the range of −1.0 and −2.2 μm.

#### Cryo-EM data processing and model building

Data were processed using CryoSPARC (Punjani et al., 2017). Movies were motion-corrected using patch motion correcting and contrast transfer function (CTF) parameters were estimated using patch CTF estimation. Micrographs were then curated to exclude those with CTF resolution below 4.5 Å, resulting in 8653 micrographs used. A 3-dimensional model from a small dataset collected previously on a Talos Arctica was used to make 30 2-dimensional templates that were used for template-based picking in CryoSPARC. 1822048 initial particles were picked. Particles were selected using three rounds of 2-D classification resulting in 452065 particles. Four ab-initio models were produced, then three rounds heterogeneous refinement were used for further particle selection, producing a final set of 178031 particles. Non-uniform refinement performed using this particle set and the best model from heterogeneous refinement produced a map at 3.4 Å. Then local refinement was performed using a mask that excluded the constant region of the 11G2 Fab, resulting in a map at 3.3 Å resolution. The map was post-processed with DeepEMhancer using the highRes model (Sanchez-Garcia et al., 2021). Atomic coordinates for ADAM10 (PDB 6BE6), 11G2 Fab (PDB 6BDZ) and the predicted structure for Tspan15 (Alphafold database O95858) were fit into the density using Chimera (Pettersen et al., 2004). Due to the structural reorganization of ADAM10, the coordinates (PDB 6BE6) were divided into two parts for fitting: the metalloproteinase domain and the disintegrin-cysteine-rich region. The model was built in Coot (Emsley and Cowtan, 2004) and refined in Phenix Real-Space Refine (Afonine et al., 2012) using secondary structure restraints. Model to map FSC was produced using Phenix Comprehensive Validation.

#### Co-immunoprecipitation and western blotting

HEK293T cells were grown in 6 well tissue culture dishes in DMEM with 10% FBS and transfected with 2 μg DNA per well (1 μg each of ADAM10-myc and FLAG-Tspan15 plasmids) using Fugene HD (Promega) according to the manufacturer’s instructions. Immunoprecipitation and western blotting were performed as described previously (Lipper et al., 2022). Primary antibodies used for western blotting were anti-FLAG (Cell Signaling Technology; rabbit, 1:2000 dilution), anti-myc (Cell Signaling Technology; rabbit, 1:2000 dilution), and anti-GAPDH (Cell Signaling Technology; rabbit, 1:2000 dilution). Anti-rabbit-HRP (Abcam) was used for the secondary antibody at 1:10000. Two biological replicates were performed.

#### N-cadherin cleavage assay

U251 Tspan15 knockout cells were described previously (Lipper et al., 2022). U251 wild-type cells and Tspan15 knockout cells were grown in 6 well tissue culture dishes in DMEM with 10% FBS. Cells were transfected with 2 μg total DNA (2 μg Ncad-HA plasmid for Figure 3; 1 ug each Ncad-HA and FLAG-Tspan plasmids for Figures 4 and 5) using 3.2 μl U251 Avalanche transfection reagent (EZ Biosystems) according to the manufacturer’s protocol. Following transfection cells were treated and subjected to western blotting as described previously (Lipper et al., 2022). Primary antibodies used were anti-HA antibody (Cell Signaling Technology; mouse, 1:1000 dilution), anti-FLAG (Cell Signaling Technology; rabbit, 1:2000 dilution), anti-β-tubulin (Cell Signaling Technology; mouse, 1:1000 dilution), or GAPDH (Cell Signaling Technology; rabbit, 1:5000). Anti-rabbit-HRP (Abcam; 1:10000 dilution) and anti-mouse-HRP (ThermoFisher; 1:10000 dilution) were used for the secondary antibodies. Blots were quantified using ImageJ software and analyzed with Graphpad Prism 7. Three biological replicates were performed.

### Supplementary Video S1

Video S1. Morph movie of the ADAM10 ectodomain converting from its closed (PDB: 6BE6) to its open, Tspan15-bound form. Related to Figure 1.

## References

Afonine, P.V., Grosse-Kunstleve, R.W., Echols, N., Headd, J.J., Moriarty, N.W., Mustyakimov, M., Terwilliger, T.C., Urzhumtsev, A., Zwart, P.H., and Adams, P.D. (2012). Towards automated crystallographic structure refinement with phenix.refine. Acta Crystallogr. D Biol. Crystallogr. 68, 352–367.

Anders, A., Gilbert, S., Garten, W., Postina, R., and Fahrenholz, F. (2001). Regulation of the α-secretase ADAM10 by its prodomain and proprotein convertases. FASEB J. 15, 1837–1839.

Dornier, E., Coumailleau, F., Ottavi, J.-F., Moretti, J., Boucheix, C., Mauduit, P., Schweisguth, F., and Rubinstein, E. (2012). TspanC8 tetraspanins regulate ADAM10/Kuzbanian trafficking and promote Notch activation in flies and mammals. J. Cell Biol. 199, 481–496.

Emsley, P., and Cowtan, K. (2004). Coot: model-building tools for molecular graphics. Acta Crystallogr. D Biol. Crystallogr. 60, 2126–2132.

Eschenbrenner, E., Jouannet, S., Clay, D., Chaker, J., Boucheix, C., Brou, C., Tomlinson, M.G., Charrin, S., and Rubinstein, E. (2020). TspanC8 tetraspanins differentially regulate ADAM10 endocytosis and half-life. Life Sci. Alliance 3, e201900444.

Esler, W.P., and Wolfe, M.S. (2001). A Portrait of Alzheimer Secretases--New Features and Familiar Faces. Science. 293, 1449–1454.

Haining, E.J., Yang, J., Bailey, R.L., Khan, K., Collier, R., Tsai, S., Watson, S.P., Frampton, J., Garcia, P., and Tomlinson, M.G. (2012). The TspanC8 Subgroup of Tetraspanins Interacts with A Disintegrin and Metalloprotease 10 (ADAM10) and Regulates Its Maturation and Cell Surface Expression*. J. Biol. Chem. 287, 39753–39765.

Harrison, N., Koo, C.Z., and Tomlinson, M.G. (2021). Regulation of ADAM10 by the TspanC8 Family of Tetraspanins and Their Therapeutic Potential. Int. J. Mol. Sci. 22, 6707.

Hartmann, D., de Strooper, B., Serneels, L., Craessaerts, K., Herreman, A., Annaert, W., Umans, L., Lübke, T., Lena Illert, A., von Figura, K., et al. (2002). The disintegrin/metalloprotease ADAM 10 is essential for Notch signalling but not for α-secretase activity in fibroblasts. Hum. Mol. Genet. 11, 2615–2624.

Hemler, M.E. (2005). Tetraspanin functions and associated microdomains. Nat. Rev. Mol. Cell Biol. 6, 801–811.

Hiroshima, K., Shiiba, M., Oka, N., Hayashi, F., Ishida, S., Fukushima, R., Koike, K., Iyoda, M., Nakashima, D., Tanzawa, H., et al. (2019). Tspan15 plays a crucial role in metastasis in oral squamous cell carcinoma. Exp. Cell Res. 384, 111622.

Jouannet, S., Saint-Pol, J., Fernandez, L., Nguyen, V., Charrin, S., Boucheix, C., Brou, C., Milhiet, P.-E., and Rubinstein, E. (2016). TspanC8 tetraspanins differentially regulate the cleavage of ADAM10 substrates, Notch activation and ADAM10 membrane compartmentalization. Cell. Mol. Life Sci. 73, 1895–1915.

Jumper, J., Evans, R., Pritzel, A., Green, T., Figurnov, M., Ronneberger, O., Tunyasuvunakool, K., Bates, R., Žídek, A., Potapenko, A., et al. (2021). Highly accurate protein structure prediction with AlphaFold. Nature 596, 583–589.

Kohutek, Z.A., diPierro, C.G., Redpath, G.T., and Hussaini, I.M. (2009). ADAM-10-Mediated N-Cadherin Cleavage Is Protein Kinase C-Dependent and Promotes Glioblastoma Cell Migration. J. Neurosci. 29, 4605–4615.

Koo, C.Z., Harrison, N., Noy, P.J., Szyroka, J., Matthews, A.L., Hsia, H.-E., Müller, S.A., Tüshaus, J., Goulding, J., Willis, K., et al. (2020). The tetraspanin Tspan15 is an essential subunit of an ADAM10 scissor complex. J. Biol. Chem. 295, 12822–12839.

Koo, C.Z., Matthews, A.L., Harrison, N., Szyroka, J., Nieswandt, B., Gardiner, E.E., Poulter, N.S., and Tomlinson, M.G. (2022). The Platelet Collagen Receptor GPVI Is Cleaved by Tspan15/ADAM10 and Tspan33/ADAM10 Molecular Scissors. Int. J. Mol. Sci. 23, 2440.

Kuhn, P.-H., Wang, H., Dislich, B., Colombo, A., Zeitschel, U., Ellwart, J.W., Kremmer, E., Roßner, S., and Lichtenthaler, S.F. (2010). ADAM10 is the physiologically relevant, constitutive α-secretase of the amyloid precursor protein in primary neurons. EMBO J. 29, 3020–3032.

Kuhn, P.-H., Colombo, A.V., Schusser, B., Dreymueller, D., Wetzel, S., Schepers, U., Herber, J., Ludwig, A., Kremmer, E., Montag, D., et al. (2016). Systematic substrate identification indicates a central role for the metalloprotease ADAM10 in axon targeting and synapse function. ELife 5, e12748.

Lambrecht, B.N., Vanderkerken, M., and Hammad, H. (2018). The emerging role of ADAM metalloproteinases in immunity. Nat. Rev. Immunol. 18, 745–758.

Lichtenthaler, S.F., Lemberg, M.K., and Fluhrer, R. (2018). Proteolytic ectodomain shedding of membrane proteins in mammals—hardware, concepts, and recent developments. EMBO J. 37 e99456.

Lipper, C.H., Gabriel, K.-H., Seegar, T.C.M., Dürr, K.L., Tomlinson, M.G., and Blacklow, S.C. (2022). Crystal structure of the Tspan15 LEL domain reveals a conserved ADAM10 binding site. Structure 30, 206–214.e4.

Liu, C., Xu, P., Lamouille, S., Xu, J., and Derynck, R. (2009). TACE-mediated ectodomain shedding of the type I TGF-beta receptor downregulates TGF-beta signaling. Mol. Cell 35, 26–36.

Liu, P.C.C., Liu, X., Li, Y., Covington, M., Wynn, R., Huber, R., Hillman, M., Yang, G., Ellis, D., Marando, C., et al. (2006). Identification of ADAM10 as a major source of HER2 ectodomain sheddase activity in HER2 overexpressing breast cancer cells. Cancer Biol. Ther. 5, 657–664.

Loganathan, S.K., Schleicher, K., Malik, A., Quevedo, R., Langille, E., Teng, K., Oh, R.H., Rathod, B., Tsai, R., Samavarchi-Tehrani, P., et al. (2020). Rare driver mutations in head and neck squamous cell carcinomas converge on NOTCH signaling. Science 367, 1264–1269.

Maretzky, T., Reiss, K., Ludwig, A., Buchholz, J., Scholz, F., Proksch, E., Strooper, B. de, Hartmann, D., and Saftig, P. (2005). ADAM10 mediates E-cadherin shedding and regulates epithelial cell-cell adhesion, migration, and β-catenin translocation. Proc. Natl. Acad. Sci. 102, 9182–9187.

Mastronarde, D.N. (2005). Automated electron microscope tomography using robust prediction of specimen movements. J. Struct. Biol. 152, 36–51.

Morin, A., Eisenbraun, B., Key, J., Sanschagrin, P.C., Timony, M.A., Ottaviano, M., and Sliz, P. (2013). Collaboration gets the most out of software. ELife 2, e01456.

Mumm, J.S., Schroeter, E.H., Saxena, M.T., Griesemer, A., Tian, X., Pan, D.J., Ray, W.J., and Kopan, R. (2000). A ligand-induced extracellular cleavage regulates gamma-secretase-like proteolytic activation of Notch1. Mol. Cell 5, 197–206.

Noy, P.J., Yang, J., Reyat, J.S., Matthews, A.L., Charlton, A.E., Furmston, J., Rogers, D.A., Rainger, G.E., and Tomlinson, M.G. (2016a). TspanC8 Tetraspanins and A Disintegrin and Metalloprotease 10 (ADAM10) Interact via Their Extracellular Regions: EVIDENCE FOR DISTINCT BINDING MECHANISMS FOR DIFFERENT TspanC8 PROTEINS. J. Biol. Chem. 291, 3145–3157.

Noy, P.J., Yang, J., Reyat, J.S., Matthews, A.L., Charlton, A.E., Furmston, J., Rogers, D.A., Rainger, G.E., and Tomlinson, M.G. (2016b). TspanC8 Tetraspanins and A Disintegrin and Metalloprotease 10 (ADAM10) Interact via Their Extracellular Regions. J. Biol. Chem. 291, 3145–3157.

Peschon, J.J., Slack, J.L., Reddy, P., Stocking, K.L., Sunnarborg, S.W., Lee, D.C., Russell, W.E., Castner, B.J., Johnson, R.S., Fitzner, J.N., et al. (1998). An Essential Role for Ectodomain Shedding in Mammalian Development. Science 282, 1281–1284.

Pettersen, E.F., Goddard, T.D., Huang, C.C., Couch, G.S., Greenblatt, D.M., Meng, E.C., and Ferrin, T.E. (2004). UCSF Chimera--a visualization system for exploratory research and analysis. J. Comput. Chem. 25, 1605–1612.

Prox, J., Willenbrock, M., Weber, S., Lehmann, T., Schmidt-Arras, D., Schwanbeck, R., Saftig, P., and Schwake, M. (2012). Tetraspanin15 regulates cellular trafficking and activity of the ectodomain sheddase ADAM10. Cell. Mol. Life Sci. 69, 2919–2932.

Pruessmeyer, J., and Ludwig, A. (2009). The good, the bad and the ugly substrates for ADAM10 and ADAM17 in brain pathology, inflammation and cancer. Semin. Cell Dev. Biol. 20, 164–174.

Punjani, A., Rubinstein, J.L., Fleet, D.J., and Brubaker, M.A. (2017). cryoSPARC: algorithms for rapid unsupervised cryo-EM structure determination. Nat. Methods 14, 290–296.

Sanchez-Garcia, R., Gomez-Blanco, J., Cuervo, A., Carazo, J.M., Sorzano, C.O.S., and Vargas, J. (2021). DeepEMhancer: a deep learning solution for cryo-EM volume post-processing. Commun. Biol. 4, 1–8.

Schrodinger, L. (2010). The PyMOL Molecular Graphics System, Version 2.5.1.

Seegar, T.C.M., Killingsworth, L.B., Saha, N., Meyer, P.A., Patra, D., Zimmerman, B., Janes, P.W., Rubinstein, E., Nikolov, D.B., Skiniotis, G., et al. (2017). Structural Basis for Regulated Proteolysis by the α-Secretase ADAM10. Cell 171, 1638–1648.e7.

Seipold, L., Altmeppen, H., Koudelka, T., Tholey, A., Kasparek, P., Sedlacek, R., Schweizer, M., Bär, J., Mikhaylova, M., Glatzel, M., et al. (2018). In vivo regulation of the A disintegrin and metalloproteinase 10 (ADAM10) by the tetraspanin 15. Cell. Mol. Life Sci. 75, 3251–3267.

Shah, J., Rouaud, F., Guerrera, D., Vasileva, E., Popov, L.M., Kelley, W.L., Rubinstein, E., Carette, J.E., Amieva, M.R., and Citi, S. (2018). A Dock-and-Lock Mechanism Clusters ADAM10 at Cell-Cell Junctions to Promote α-Toxin Cytotoxicity. Cell Rep. 25, 2132–2147.e7.

Solanas, G., Cortina, C., Sevillano, M., and Batlle, E. (2011). Cleavage of E-cadherin by ADAM10 mediates epithelial cell sorting downstream of EphB signalling. Nat. Cell Biol. 13, 1100–1107.

Sprinzak, D., and Blacklow, S.C. (2021). Biophysics of Notch Signaling. Annu. Rev. Biophys 50, 157–189.

Suh, J., Choi, S.H., Romano, D.M., Gannon, M.A., Lesinski, A.N., Kim, D.Y., and Tanzi, R.E. (2013). ADAM10 missense mutations potentiate β-amyloid accumulation by impairing prodomain chaperone function. Neuron 80, 385–401.

Sulis, M.L., Saftig, P., and Ferrando, A.A. (2011). Redundancy and specificity of the metalloprotease system mediating oncogenic NOTCH1 activation in T-ALL. Leukemia 25, 1564–1569.

Susa, K.J., Seegar, T.C., Blacklow, S.C., and Kruse, A.C. (2020). A dynamic interaction between CD19 and the tetraspanin CD81 controls B cell co-receptor trafficking. ELife 9, e52337.

Susa, K.J., Rawson, S., Kruse, A.C., and Blacklow, S.C. (2021). Cryo-EM structure of the B cell coreceptor CD19 bound to the tetraspanin CD81. Science 371, 300–305.

Uemura, K., Kihara, T., Kuzuya, A., Okawa, K., Nishimoto, T., Ninomiya, H., Sugimoto, H., Kinoshita, A., and Shimohama, S. (2006). Characterization of sequential N-cadherin cleavage by ADAM10 and PS1. Neurosci. Lett. 402, 278–283.

Umeda, R., Satouh, Y., Takemoto, M., Nakada-Nakura, Y., Liu, K., Yokoyama, T., Shirouzu, M., Iwata, S., Nomura, N., Sato, K., et al. (2020). Structural insights into tetraspanin CD9 function. Nat. Commun. 11, 1606.

Venkatesh, H.S., Tam, L.T., Woo, P.J., Lennon, J., Nagaraja, S., Gillespie, S.M., Ni, J., Duveau, D.Y., Morris, P.J., Zhao, J.J., et al. (2017). Targeting neuronal activity-regulated neuroligin-3 dependency in high-grade glioma. Nature 549, 533–537.

Weng, A.P., Ferrando, A.A., Lee, W., Morris, J.P., Silverman, L.B., Sanchez-Irizarry, C., Blacklow, S.C., Look, A.T., and Aster, J.C. (2004). Activating mutations of NOTCH1 in human T cell acute lymphoblastic leukemia. Science 306, 269–271.

Weskamp, G., Ford, J.W., Sturgill, J., Martin, S., Docherty, A.J.P., Swendeman, S., Broadway, N., Hartmann, D., Saftig, P., Umland, S., et al. (2006). ADAM10 is a principal “sheddase” of the low-affinity immunoglobulin E receptor CD23. Nat. Immunol. 7, 1293–1298.

Wetzel, S., Seipold, L., and Saftig, P. (2017). The metalloproteinase ADAM10: A useful therapeutic target? Biochim. Biophys. Acta BBA - Mol. Cell Res. 1864, 2071–2081.

Yang, Y., Liu, X.R., Greenberg, Z.J., Zhou, F., He, P., Fan, L., Liu, S., Shen, G., Egawa, T., Gross, M.L., et al. (2020). Open conformation of tetraspanins shapes interaction partner networks on cell membranes. EMBO J. 39, e105246.

Zhang, B., Zhang, Z., Li, L., Qin, Y.-R., Liu, H., Jiang, C., Zeng, T.-T., Li, M.-Q., Xie, D., Li, Y., et al. (2018). TSPAN15 interacts with BTRC to promote oesophageal squamous cell carcinoma metastasis via activating NF-κB signaling. Nat. Commun. 9, 1423.

Zimmerman, B., Kelly, B., McMillan, B.J., Seegar, T.C.M., Dror, R.O., Kruse, A.C., and Blacklow, S.C. (2016). Crystal Structure of a Full-Length Human Tetraspanin Reveals a Cholesterol-Binding Pocket. Cell 167, 1041–1051.e11.

